# LazyAF, a pipeline for accessible medium-scale *in silico* prediction of protein-protein interactions

**DOI:** 10.1101/2024.01.29.577767

**Authors:** Thomas C. McLean

## Abstract

Artificial intelligence has revolutionized the field of protein structure prediction. However, with more powerful and complex software being developed, it is accessibility and ease of use rather than capability that is quickly becoming a limiting factor to end users. Here, I present a Google Colaboratory-based pipeline, named LazyAF, which integrates the existing ColabFold BATCH to streamline the process of medium-scale protein-protein interaction prediction. I apply LazyAF to predict the interactome of the 76 proteins encoded on a broad-host-range multi-drug resistance plasmid RK2, demonstrating the ease and accessibility the pipeline provides.

**Availability:** LazyAF is freely available at https://github.com/ThomasCMcLean/LazyAF

## INTRODUCTION

The integration of artificial intelligence-based protein structure prediction software, such as AlphaFold2 and RoseTTAFold, into hosted notebook services such as Google Colaboratory has dramatically improved accessibility (Baek et al., 2021; Jumper et al., 2021; Mirdita et al., 2022). These platforms are not only a user-friendly but also provide access to powerful graphics processing units (GPU) to accelerate predictions of large or complex protein structures, thereby removing the requirement for local high-performance computing clusters that are not widely available or require advanced computer literacy to use.

As the speed and accessibility of protein structure prediction increase so does the potential scale, ranging from single protein structure predictions to screening large datasets of proteins for new protein–protein interactions (PPI). Here, I designed a Google Colaboratory-based LazyAF pipeline to work with ColabFold BATCH (Mirdita et al., 2022) to streamline the process of screening large datasets of proteins for novel PPIs and to make it accessible to users with less bioinformatics experience. I demonstrate the accessibility of this pipeline and its ability to perform automatic medium-throughput prediction of protein complexes in an all-versus-all scenario using the 76 proteins encoded on a multidrug-resistance plasmid RK2 as an example (Pansegrau et al., 1994). With LazyAF, I hope to lower the entry barrier for medium-scale PPI prediction and empower more wet-bench researchers to integrate *in silico* modeling into their workflow.

## SOFTWARE DESCRIPTION

AlphaFold2-Multimer-based prediction of PPIs is conceptually similar to a co-immunoprecipitation experiment where a protein of interest is used as bait to pull down its interacting protein partners from the total cell lysate. LazyAF takes a protein as a ‘bait’ and a list of other proteins as ‘candidates’ to automatically run an AlphaFold2-Multimer prediction between each bait and candidate before ranking the likelihood of their interactions. LazyAF consists of two open-source Google Colaboratory notebooks (see data availability). At the core of LazyAF is ColabFold BATCH which is also available via Google Colaboratory (https://github.com/sokrypton/ColabFold) (Kim et al., 2023; Mirdita et al., 2022). An outline of LazyAF is depicted in Figure 1. A step-by-step protocol can be found in Supplementary Note 1.

**Figure 1.**
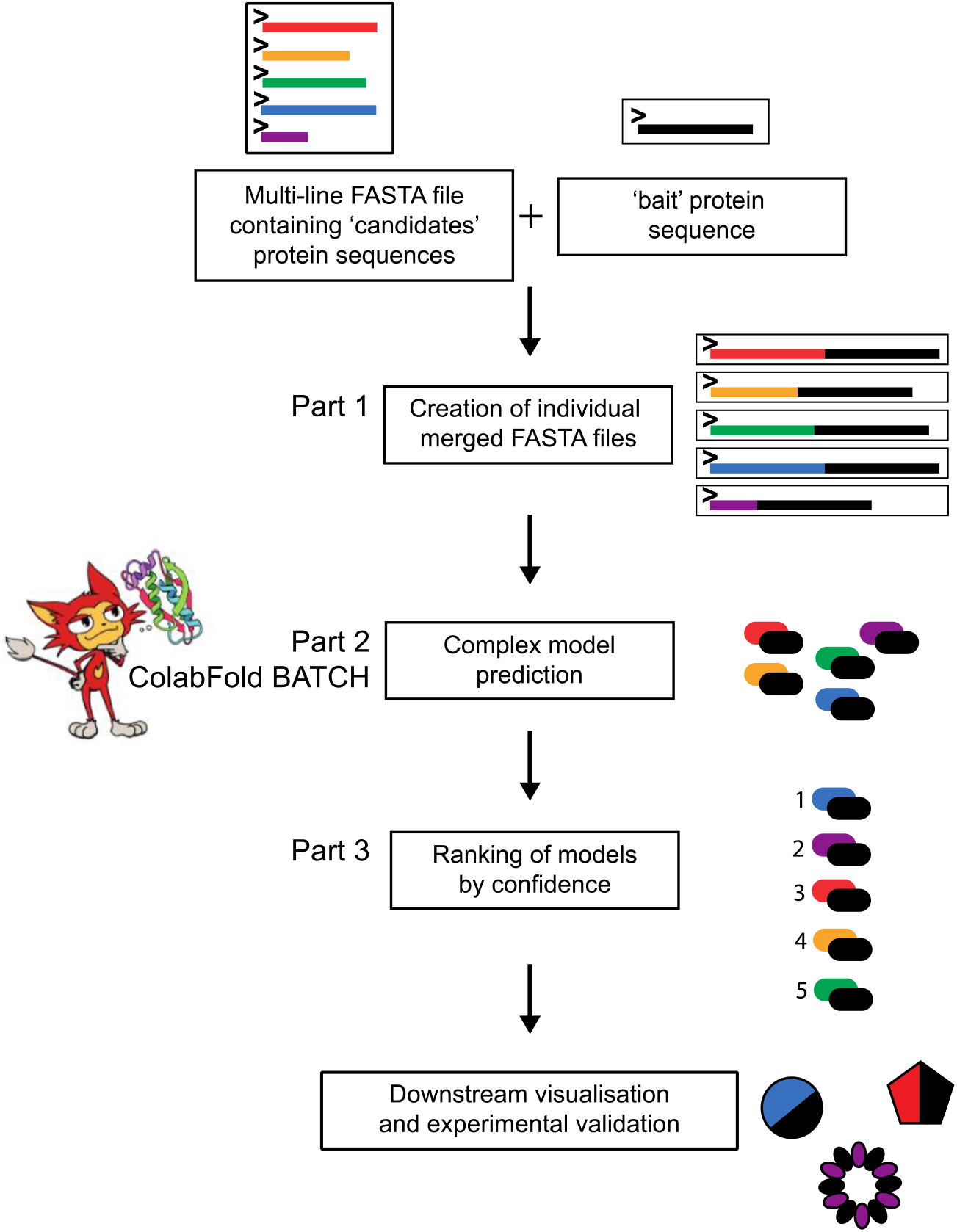
The LazyAF pipeline. LazyAF Part 1 concatenates a FASTA file containing multiple ‘candidate’ protein sequences with a single ‘bait’ protein sequence and outputs each as an individual FASTA file. This collection of individual FASTA files is then used as the input for AlphaFold2-Multimer-based protein structure prediction using ColabFold BATCH. This takes advantage of powerful cloud-based GPUs to enable rapid modelling without the need of local high-performance computing facilities. LazyAF Part 3 retrieves the results from the modeling in Part 2 and ranks the likelihood of protein-protein interactions by their ranking confidence score. These results can be used to generate new hypotheses but should always be experimentally validated.

### Part 1 – preparation of input files for ColabFold BATCH

Input for LazyAF Part 1 consists of (i) a name and sequence of a ‘bait’ protein and (ii) a multi-line FASTA file containing sequences of ‘candidate’ proteins. It is recommended to download the ‘candidate’ sequence file from NCBI (see Supplementary Note 1) as this file contains a unique identifier for each candidate protein sequence.

The output of LazyAF Part 1 is a collection of individual FASTA files each with the sequence of the ‘bait’ and that of a ‘candidate’ joined via a colon. A colon separator designates separate chains for the subsequent AlphaFold2-Multimer-based protein co-folding. Each FASTA file is uniquely named based on the name of the ‘bait’ and the unique identifier of each ‘candidate’ protein. All individual FASTA files are stored in a cloud-based Google Drive folder.

### Part 2 – structure prediction in a batch mode by ColabFold BATCH

The output folder from LazyAF Part 1 can be used directly as the input for ColabFold BATCH. For further information and methodology on ColabFold BATCH see Mirdita et al., 2022.

### Part 3 – ranking likelihood of bait-candidate interactions

LazyAF Part 3 simplifies the collation and analysis of the numerous PPI predictions from ColabFold BATCH (Mirdita et al., 2022). With each prediction producing five models alongside other accessory files, the amount of data is large and challenging to analyze manually. LazyAF Part 3 locates the JSON file associated with the top-ranked prediction for each co-folding and copies them into a new folder for subsequent analysis. The predicted template modeling score (pTM) and predicted interface template modeling score (ipTM) are then extracted from these JSON files, and a ranking score (ranking confidence) is calculated for each PPI using the following formula (ranking confidence = 0.2pTM + 0.8ipTM) (Wallner, 2023a). These ranking confidence scores can then be sorted from high to low to assess the quality of models and the likelihood of interaction between each bait-candidate protein pair.

### LazyAF predicts the potential interactome of 76 proteins encoded on the RK2 plasmid

I demonstrate the accessibility of LazyAF and its ability to perform automatic medium-throughput prediction of protein complexes in an all-versus-all scenario using 76 proteins encoded on a multi-drug resistance plasmid RK2. A multi-line FASTA file containing all protein sequences encoded on the RK2 genome (BN000925.1) was retrieved from NCBI and uploaded to Google Drive. Each protein sequence (76 in total) was used as a bait sequence in LazyAF Part 1 to generate 76x76 i.e. 5,776 FASTA files with every possible pairwise combination of protein-protein interactions. This folder was then used as the input directory for ColabFold v1.5.2 AlphaFold2 MMseqs2 BATCH with the following settings: msa_mode: MMseqs2 (UniRef+Environmental), num_models: 5, num_recycles: 3, stop_at_score: 100. The notebook was run on High-RAM A100 or V100 GPUs depending on their availability on Google Colaboratory. The resulting folder containing the outputs from AlphaFold2 predictions was used as the input directory for LazyAF Part 3 which copied the top-ranked JSON files for each predicted PPI into an analysis folder. Part 3 also retrieved the pTM and ipTM scores for each top-ranked model and calculated the ranking confidence score (0.2pTM + 0.8ipTM). These values were stored in an output CSV file on Google Drive (Supplementary Table 1). A heatmap of ranking confidence scores for 5,776 PPIs was plotted using GraphPad Prism 10 (Figure 2), and an interaction network was generated using Cytoscape (Figure 3).

**Figure 2.**
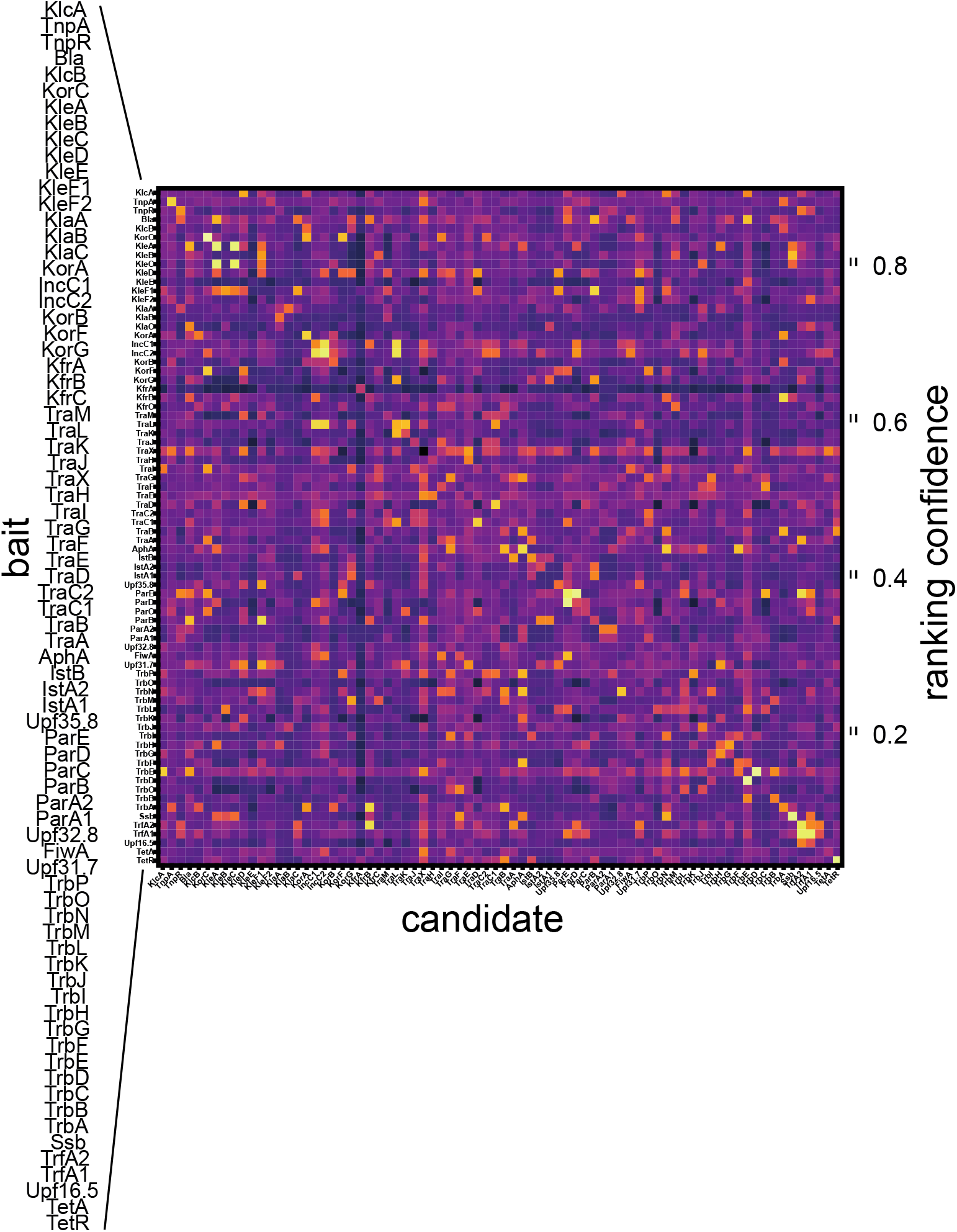
Heatmap displaying ranking confidence scores for 5,776 interactions from the 76 proteins encoded on the RK2 plasmid. Proteins are ordered by the sequential genomic locations of their encoding genes on RK2. The ranking confidence score is calculated using the formula: ranking confidence score = 0.2pTM + 0.8ipTM for the top ranked model of each protein-protein pair prediction.

**Figure 3.**
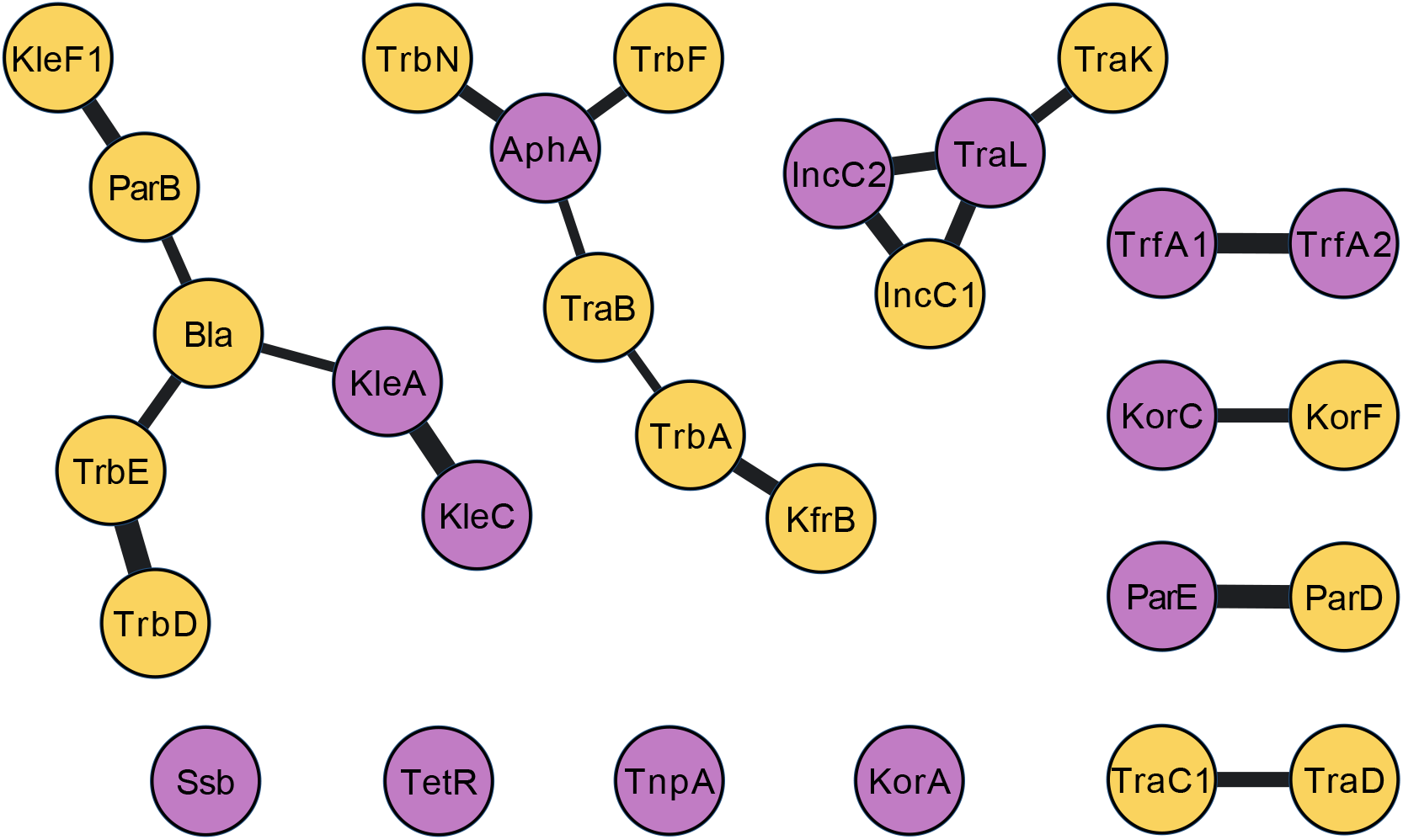
An interaction network for most likely protein-protein interaction pair from the all-versus-all screen of 5,776 potential interactions from the 76 proteins encoded on the RK2 plasmid. Yellow nodes indication proteins predicted to not self-interact. Purple nodes indicate proteins predicted to self-interact. Nodes are included if their ranking confidence scores are > 0.7 in both primary and reciprocal pairings. The thickness of the line is proportional to a maximal ranking confidence score.

I noted that the heatmap of ranking confidence scores is not symmetrical (Figure 2), suggesting that a score of co-folding between protein A (bait):protein B (candidate) sometimes is different from that of a co-folding between protein B (bait):protein A (candidate). Variation is to be expected due to the nature of model building by iterative refinement, however to reduce the likelihood of false positives I only considered the PPIs with a ranking confidence score of > 0.7 in both directions (O’Reilly et al., 2023; Wallner, 2023b). Among 5,776 predictions, 50 predicted PPIs have ranking confidence scores > 0.7 in a single direction (i.e. protein A (bait):protein B (candidate)) and also in the reciprocal direction (i.e. protein B (bait):protein A (candidate)). I consider these 50 predicted PPIs to be of high likelihood (Figure 3). Several of the top predicted PPIs have been experimentally validated previously, particularly the homodimerization of transcriptional regulators KorA (Jagura-Burdzy and Thomas, 1995), TetR (Ramos Juan L. et al., 2005), and the transposase TnpA (Ronning et al., 2005). Our screen also suggested many new potential protein complexes. These should be experimentally validated and characterized to further understand the biology of the broad-host-range RK2 plasmid.

## CONCLUSION

LazyAF provides a highly accessible pipeline that can be used in any web browser and can utilize powerful cloud-based hardware to facilitate predictions. Due diligence should always be taken to manually examine models, checking other parameters such as the per-residue confidence (pLDDT), the pDockQ/mpDockQ scores (Bryant et al., 2022b, 2022a), and validation by complementary *in vitro*/*in vivo* methodologies. Alternative software, such as AlphaPulldown (Yu et al., 2023), exists and goes well beyond the capabilities of LazyAF. However, this Python-based package requires substantial bioinformatic expertise to install and execute. LazyAF provides a slimmed-down, accessible pipeline for *in silico* pulldown experiments to users with less bioinformatics experience. As the line blurs between traditional wet and dry laboratory research, I anticipate that LazyAF will be of most use to wet-lab scientists who are keen to integrate *in silico* protein structure predictions into their workflow.

## Acknowledgments

This toolkit is built around the great work of Sergey Ovchinnikov (@sokrypton), Milot Mirdita (@milot_mirdita) and Martin Steinegger (@thesteinegger) (Mirdita et al., 2022). I thank Tung Le for his invaluable mentoring, funding and, most importantly, his support for this project. I thank Andres Posbeyikian and Sophien Kamoun for their detailed introduction to Colabfold BATCH and the instrumental tips and tricks they provided. I thank Patrick Allen for helpful comments on the pipeline. I also thank Emma Banks, Rebecca Diss, Leah McPhillips, and Max Jordan for testing the pipeline and for useful suggestions to improve usability.

## SUPPLEMENTARY INFORMATION

**Supplementary Table 1.** Output from LazyAF Part 3 for PPIs with ranking confidence scores > 0.7.

| Candidate | Bait | pTM | ipTM | Ranking_confidence |
| --- | --- | --- | --- | --- |
| KorC | KorC | 0.91 | 0.89 | 0.894 |
| TrbE | TrbD | 0.85 | 0.89 | 0.882 |
| TrbD | TrbE | 0.85 | 0.89 | 0.882 |
| ParE | ParD | 0.77 | 0.9 | 0.874 |
| ParD | ParE | 0.76 | 0.9 | 0.872 |
| Ssb | Ssb | 0.87 | 0.87 | 0.87 |
| TetR | TetR | 0.87 | 0.87 | 0.87 |
| KleA | KleA | 0.86 | 0.86 | 0.86 |
| KleA | KleC | 0.86 | 0.86 | 0.86 |
| IncC2 | IncC2 | 0.9 | 0.84 | 0.852 |
| KleC | KleA | 0.86 | 0.85 | 0.852 |
| TrfA1 | TrfA1 | 0.78 | 0.86 | 0.844 |
| TrfA1 | TrfA2 | 0.78 | 0.86 | 0.844 |
| KleC | KleC | 0.85 | 0.84 | 0.842 |
| ParE | ParE | 0.88 | 0.83 | 0.84 |
| TrfA2 | TrfA2 | 0.86 | 0.82 | 0.828 |
| IncC2 | TraL | 0.88 | 0.81 | 0.824 |
| IncC2 | IncC1 | 0.77 | 0.83 | 0.818 |
| IncC1 | IncC2 | 0.77 | 0.83 | 0.818 |
| TrfA2 | KfrB | 0.88 | 0.8 | 0.816 |
| ParE | TrfA2 | 0.87 | 0.8 | 0.814 |
| AphA | AphA | 0.85 | 0.8 | 0.81 |
| ParB | KleF1 | 0.81 | 0.81 | 0.81 |
| TraL | IncC1 | 0.76 | 0.82 | 0.808 |
| IncC1 | TraL | 0.76 | 0.82 | 0.808 |
| TrbA | KfrB | 0.82 | 0.8 | 0.804 |
| KfrB | TrbA | 0.82 | 0.8 | 0.804 |
| KleF1 | ParB | 0.81 | 0.8 | 0.802 |
| TraL | IncC2 | 0.86 | 0.78 | 0.796 |
| KorA | KorA | 0.78 | 0.8 | 0.796 |
| TnpA | TnpA | 0.82 | 0.79 | 0.796 |
| TraC1 | TraD | 0.72 | 0.81 | 0.792 |
| TrbE | KlcA | 0.79 | 0.79 | 0.79 |
| TraD | TraC1 | 0.71 | 0.81 | 0.79 |
| KorF | KorC | 0.47 | 0.86 | 0.782 |
| AphA | TrbN | 0.74 | 0.79 | 0.78 |
| TrbN | Upf32.8 | 0.71 | 0.79 | 0.774 |
| Bla | TrbE | 0.8 | 0.76 | 0.768 |
| KorC | KorF | 0.47 | 0.84 | 0.766 |
| AphA | TrbF | 0.71 | 0.78 | 0.766 |
| TrbN | AphA | 0.74 | 0.77 | 0.764 |
| KleB | Ssb | 0.81 | 0.75 | 0.762 |
| TraL | TraK | 0.69 | 0.78 | 0.762 |
| ParB | Bla | 0.84 | 0.74 | 0.76 |
| TrbF | AphA | 0.71 | 0.77 | 0.758 |
| FiwA | ParE | 0.78 | 0.75 | 0.756 |
| Bla | ParB | 0.84 | 0.73 | 0.752 |
| TraK | TraL | 0.67 | 0.77 | 0.75 |
| TrbP | AphA | 0.81 | 0.73 | 0.746 |
| Bla | KleA | 0.85 | 0.72 | 0.746 |
| TraL | TraL | 0.84 | 0.72 | 0.744 |
| AphA | TraB | 0.79 | 0.73 | 0.742 |
| Upf31.7 | KleF1 | 0.74 | 0.74 | 0.74 |
| KleA | Bla | 0.85 | 0.71 | 0.738 |
| ParE | TrbC | 0.53 | 0.79 | 0.738 |
| KorC | KorA | 0.72 | 0.74 | 0.736 |
| TrbA | TraB | 0.76 | 0.73 | 0.736 |
| TraB | TrbA | 0.76 | 0.73 | 0.736 |
| Ssb | TrbN | 0.68 | 0.75 | 0.736 |
| TraA | TrfA2 | 0.84 | 0.7 | 0.728 |
| KlcA | KleD | 0.79 | 0.71 | 0.726 |
| KorG | TraL | 0.74 | 0.72 | 0.724 |
| Upf35.8 | KleF1 | 0.68 | 0.73 | 0.72 |
| KleF1 | KlaC | 0.56 | 0.75 | 0.712 |
| TrbM | KorG | 0.66 | 0.72 | 0.708 |
| Upf31.7 | TrbH | 0.74 | 0.7 | 0.708 |
| TraB | AphA | 0.77 | 0.69 | 0.706 |
| TrbE | Bla | 0.77 | 0.69 | 0.706 |
| TraX | TraE | 0.81 | 0.68 | 0.706 |
| Tral | KorC | 0.56 | 0.74 | 0.704 |
| IstB | TraA | 0.76 | 0.69 | 0.704 |
| TrbH | TrbG | 0.63 | 0.72 | 0.702 |

### Supplementary Note 1

A step-by-step protocol on how to run LazyAF

1. Download your genome of interest from NCBI as coding sequences in the FASTA Protein format (see picture below). This will give you a file named *sequence*.*txt* containing a list of FASTAs for every coding sequence identified within the genome of interest. This will be the ‘candidate’ proteins file. I suggest renaming this *sequence*.*txt* file to include the genome name. *Note: alternatively, you can curate your version of the ‘candidate’ protein file by manually pasting protein sequences in FASTA format into a text file*.

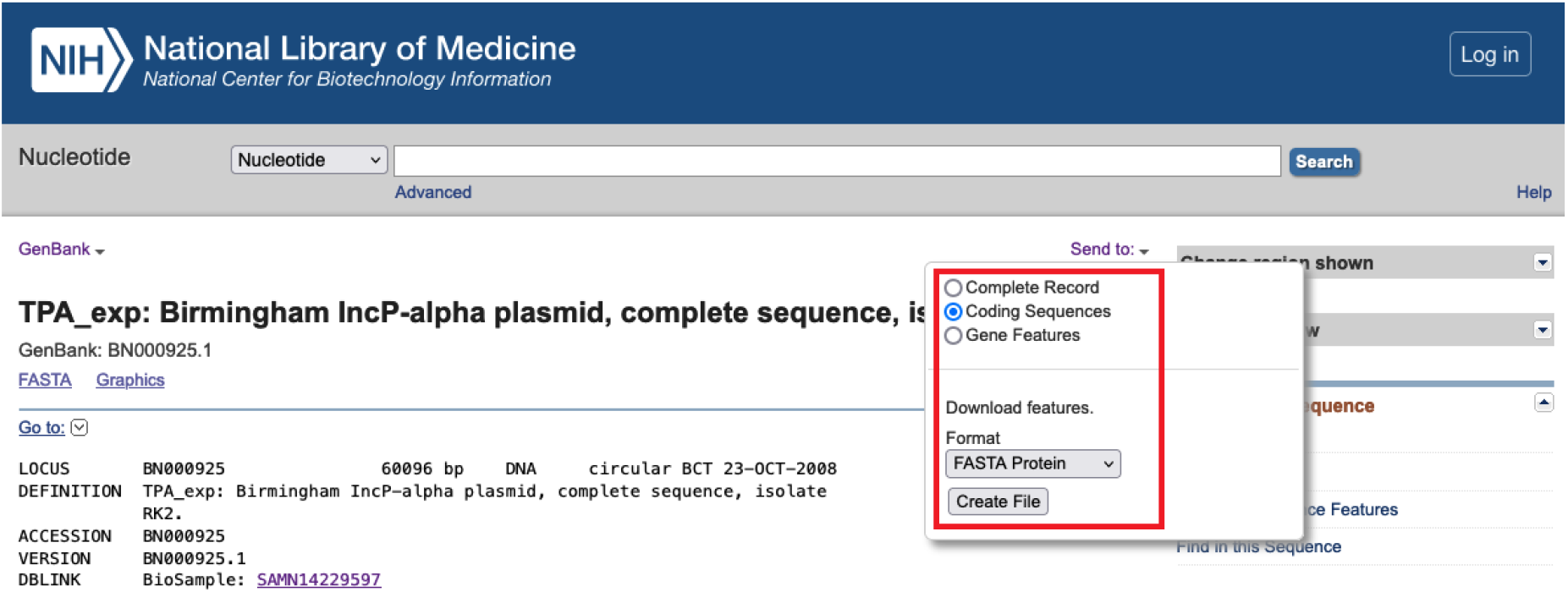
2. Create a folder in Google Drive in the *My Drive* section with the name *input* (see picture below). Upload your *sequence*.*txt* file from Step 1 to the *input* folder. *Note: you will need to register for a Google Drive account if you do not already have one*.

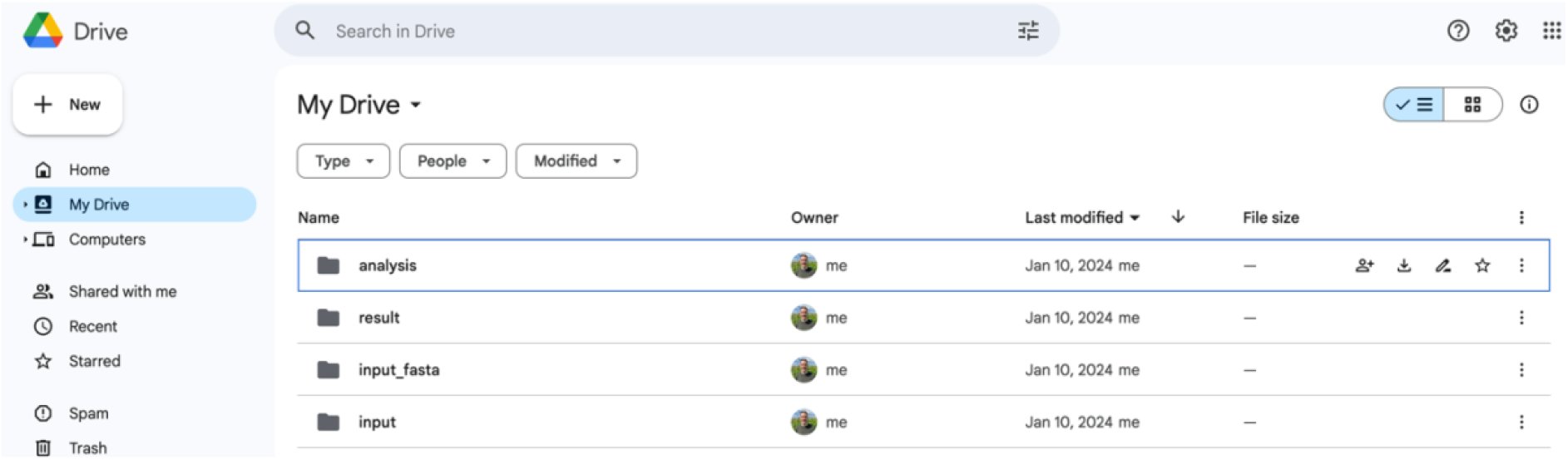
3. Open [LazyAF Part 1](https://colab.research.google.com/drive/1a5d7xraEK4Iv3Ecmmjb1opnU5jwXRW_a) in your web browser (see picture below). If you have renamed any folders or files make sure these match up to the *input_dir, result_dir* and *input_file*. Then provide the bait protein sequence and name while making sure to remove any non-amino acid letters, symbols, or spaces (a common one can be a terminal *) from the protein sequence. When ready click *runtime -> run all* to run the script. *Note: connecting to your Google Drive is required. The time required to complete the process depends on the number of coding sequences in the ‘candidate’ protein file. You can follow the process in the Google Drive input_fasta folder but on average a few minutes should suffice to complete this step*.

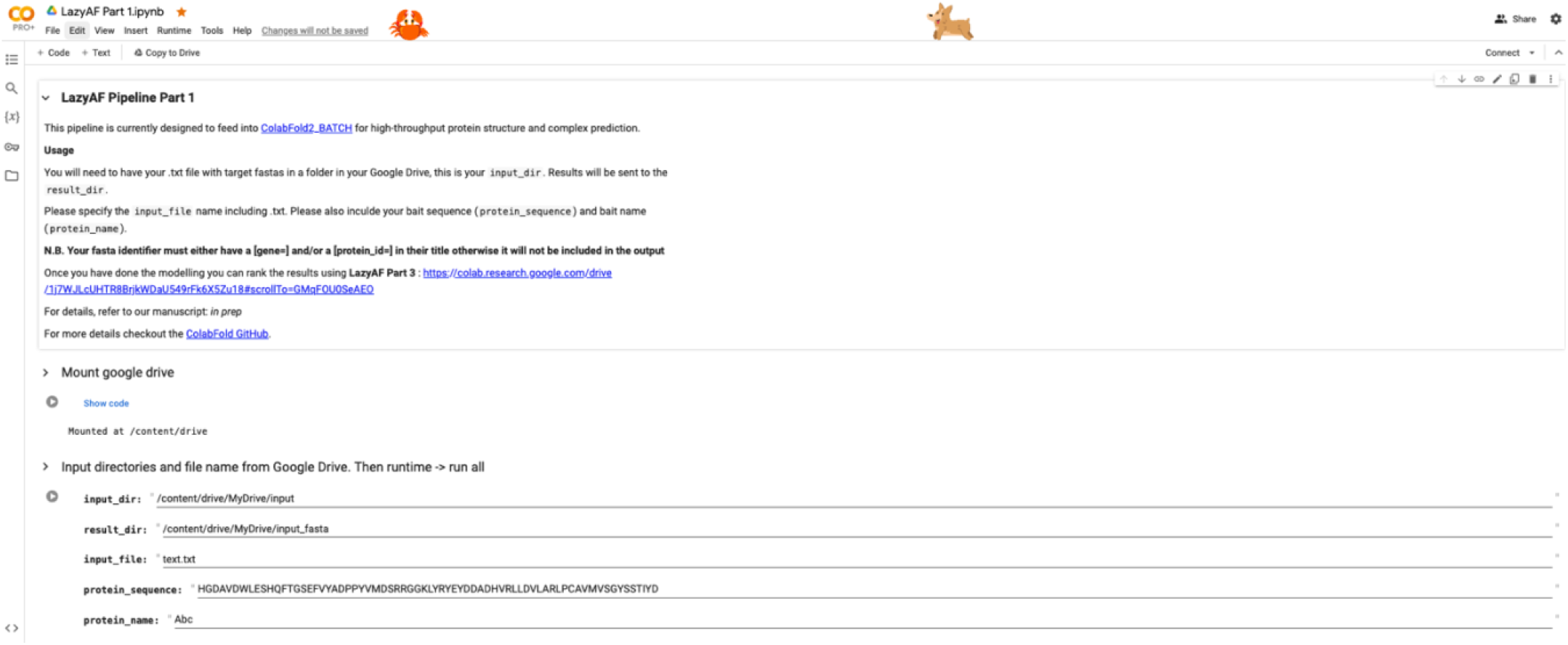
4. Once complete proceed to [LazyAF Part 2](https://colab.research.google.com/github/sokrypton/ColabFold/blob/main/batch/AlphaFold2_batch.ipynb) (Currently ColabFold v1.5.5: AlphaFold2 w/ MMseqs2 BATCH) and follow the detailed instructions provided. If you have not modified folder names then the default should match the ones in your Google Drive (N.B. there is no need to create the *result_dir* folder, the software creates it as required). For new users, I recommend using the default settings. If you have access to the Pro or Pro+ plans, you should change your GPU to either the preferable A100 or the V100 if the A100 is not available. This can be done by clicking *runtime -> change runtime type -> select A100 or V100*. When ready, click *runtime -> run all* to run the script. *Note: connecting to your Google Drive is required. The output from this modeling step can require large storage space on your Google Drive, I recommend upgrading your Google Drive plan to 100 Gb of storage capacity. I also recommend upgrading your Google Colab plan to Pro+. The upgrade not only gives users access to the more powerful hardware i*.*e. A100 GPU but also allows running of the notebook in the background. The compute unit requirements and time for complete modeling depend on the number of protein-protein interactions and the size of each protein. As a rule of thumb, if you are using an A100 GPU you will use ~15-20 compute units an hour and might model between 5-20 interactions per hour. Make sure to keep your compute units (drop-down arrow in the top right of the screen -> view resources) topped up to prevent disconnection from your runtime. ColabFold v1*.*5*.*5: AlphaFold2 w/ MMseqs2 BATCH tracks its own progress. If ColabFold is disconnected for any reason or crashes, click runtime -> disconnect and delete runtime, refresh your browser page, and restart the process. ColabFold BATCH will pick up where it left off*.
5. Once all modeling is complete proceed to [LazyAF Part 3](https://colab.research.google.com/drive/1j7WJLcUHTR8BrjkWDaU549rFk6X5Zu18). Make sure all the directories are named correctly and change the name of the output analysis file *csv_name* if you wish. Unless you have modified the naming scheme of the input files, then leave the fields *split_1* and *split_2* as default. When ready, click *runtime -> run all* to run the script. *Note: connecting to your Google Drive is required. The time required to complete the process depends on the number of predictions. You can follow the process in the Google Drive analysis folder but on average a few minutes should suffice for most jobs. When complete the folder should contain both the top-ranked JSON files for each model and also the output analysis CSV file containing columns for the bait and candidate proteins, pTM, ipTM, and ranking_confidence score*.

